# A non-Dicer RNase III and four other novel factors required for RNAi-mediated transposon suppression in the human pathogenic yeast *C. neoformans*

**DOI:** 10.1101/577924

**Authors:** Jordan E. Burke, Adam D. Longhurst, Prashanthi Natarajan, Beiduo Rao, S. John Liu, Jade Sales-Lee, Yasaman Mortensen, James J. Moresco, Jolene K. Diedrich, John R. Yates, Hiten D. Madhani

**Author notes:** **Corresponding author:** Dr. Hiten D. Madhani, N372C Genentech Hall, 600 16^th^ Street, San Francisco, CA 94158, Electronic address.

## Abstract

The human pathogenic yeast *Cryptococcus neoformans* silences transposable elements using endo-siRNAs and an Argonaute, Ago1. Endo-siRNAs production requires the RNA-dependent RNA polymerase, Rdp1, and two partially redundant Dicer enzymes, Dcr1 and Dcr2, but is independent of histone H3 lysine 9 methylation. We describe here an insertional mutagenesis screen for factors required to suppress the mobilization of the *C. neoformans HARBINGER* family DNA transposon *HAR1*. Validation experiments uncovered five novel genes (*RDE1-5*) required for *HAR1* suppression and global production of suppressive endo-siRNAs. Loss of the *RDE* genes does not impact transcript levels, suggesting the endo-siRNAs do not act by impacting target transcript synthesis or turnover. *RDE3* encodes a non-Dicer RNase III related to *S. cerevisiae* Rnt1, *RDE4* encodes a predicted terminal nucleotidyltransferase, while *RDE5* has no strongly predicted encoded domains. Affinity purification-mass spectrometry studies reveal that Rde3 and Rde5 are physically associated. *RDE1* encodes a G-patch protein homologous to the *S. cerevisiae* Sqs1/Pfa1, a nucleolar protein that directly activates the essential helicase Prp43 during rRNA biogenesis. Rde1 copurifies Rde2, another novel protein obtained in the screen, as well as Ago1, a homolog of Prp43, and numerous predicted nucleolar proteins. We also describe the isolation of conditional alleles of *PRP43*, which are defective in RNAi. This work reveals unanticipated requirements for a non-Dicer RNase III and presumptive nucleolar factors for endo-siRNA biogenesis and transposon mobilization suppression in *C. neoformans*.

## Introduction

Transposons are ancient mobile genetic elements that have invaded the genomes of nearly all living organisms and can make up substantial proportions of host DNA (Nekrutenko and Li 2001). DNA transposons and retrotransposons that retain their ability to mobilize and proliferate can both mutagenize and disrupt regulation of the host genome (Chuong *et al*. 2017). Host organisms such as fungi have therefore developed a diverse set of mechanisms to defend against transposable elements, ranging from silencing via histone modifications and DNA methylation to quelling and RNA interference (reviewed in Billmyre *et al*. 2013).

In *Cryptococcus neoformans*, transposable elements are silenced by endogenously produced small RNAs (endo-siRNAs) (Wang *et al*. 2010, 2012, 2013). *C. neoformans* harbors a semi-canonical RNAi pathway including of a single Argonaute (Ago1), two partially redundant Dicers (Dcr1, Dcr2) and an RNA-dependent RNA polymerase (Rdp1) (Janbon *et al*. 2010). Ago1 is found in both a nuclear complex, SCANR, and a complex with Gwo1 that localizes to P-bodies, the PRSC (Dumesic *et al*. 2013). siRNA biogenesis and transposon silencing appear to be particularly important during meiosis (Wang *et al*. 2010); however, transposon mobilization may also occur in vegetative cells and is suppressed by the RNAi machinery (Wang *et al*. 2010; Dumesic *et al*. 2013).

While endo-siRNA systems vary from organism to organism (Claycomb 2014), one common characteristic is the presence of a triggering double stranded RNA (dsRNA) species. In some cases, the dsRNA is produced by transcription of repetitive elements that form inter- and intra-molecular duplexes (Sijen and Plasterk 2003; Slotkin *et al*. 2005). In other organisms, an RNA-dependent RNA polymerase, such as the *C. neoformans* Rdp1, is thought to produce dsRNA (Lee *et al*. 2010; Janbon *et al*. 2010). In the latter case, the trigger for production of dsRNA is unclear, as is the manner in which the RNA-dependent RNA polymerase is recruited. More generally, how invasive genetic elements are detected by the host genome defense machinery remains unclear.

Genome defense mechanisms can be triggered by introduction of transgenes and repetitive sequences. In fact, several transposon silencing pathways were discovered due to co-suppression of transgenes and cognate endogenous genes (Billmyre *et al*. 2013). In *C. neoformans*, the RNAi pathway was initially characterized by studying the co-suppression of multiple copies of a *SXI2**a**-URA5* transgene (Wang *et al*. 2010). Upon mating and selection on media that selects against uracil biosynthesis, strains can be recovered that silence all *URA5* loci by RNAi. This suggests that cells sense either the copy number or expression level of genes and targets the RNAi pathway against anomalous transcripts. However, the manner in which such events are detected is unknown.

To pursue this question, we searched for additional components of *C. neoformans* endo-siRNA production pathway, using an active DNA transposon, *HAR1*, a member of the *HARBINGER* family as a reporter for the ability of the cell to silence transposable elements. *HARBINGER* is a cut-and-paste DNA transposon with a DDE nuclease that excises double-stranded DNA directly and reinserts the gene encoding the transposon elsewhere in the host genome (Muszewska *et al*. 2017). We took advantage of an active copy of *HARBINGER* that is silenced by the RNAi machinery in *C. neoformans* (Wang *et al*. 2010) to search for factors required for transposable element silencing. We report here five new genes required for production of endo-siRNAs and the suppression of *HAR1* mobilization.

## Materials and Methods

### Strain construction

The *ura5::HAR1:NEO* transposition screening strain was constructed using a plasmid containing the second intron of *URA5* interrupted by the *HAR1* sequence plus 1 kb of upstream flanking sequence, followed by the 3’ end of *URA5* and 500 bp of downstream flanking sequence and the *C. neoformans* G418 resistance cassette by in vivo recombination in *S. cerevisiae* into the pRS416 backbone (Finnigan and Thorner 2015). Plasmids were recovered by preparing DNA from *S. cerevisiae* and electroporation into DH5α *E. coli* followed by miniprep. The plasmid was linearized by restriction digest with PmeI and SbfI (NEB) and incorporated into the genomes of CM018 and Kn99a by biolistic transformation (Chun and Madhani 2010). Incorporation was confirmed by colony PCR and Sanger sequencing. Knockouts were incorporated into the strain by mating on Murashige and Skoog medium as previously described (Xue *et al*. 2007) followed by selection on G418 and nourseothricin (NAT). Isolates with uracil prototrophy were selected against on 5-FOA before characterization of *HARBINGER* transposition.

Proteins of interest were C-terminally tagged with CBP-2xFLAG. The tag was incorporated immediately before the annotated stop codon and followed by a terminator and the G418 resistance cassette. Linearized plasmid was introduced into Kn99a by biolistic transformation and the presence and sequence of the tag was confirmed by colony PCR, Sanger sequencing and Western blotting against the FLAG epitope.

### Transposition assay

Transposition of the *HARBINGER* transposon out of the *URA5* intron was assayed by selecting for growth on synthetic complete medium lacking uracil (SC-Ura). Strains were initially recovered from frozen stocks on YPAD, then selected for uracil auxotrophy by patching on 5-FOA plates. Enough cells to achieve 0.2-0.3 OD were resuspended in YPAD liquid medium and incubated with shaking at 30°C until doubled. 1 ml of cells were then concentrated by centrifugation at 2000xg, resuspended in 0.2 ml of supernatant and spread onto -Ura plates. Colonies were counted after 3-6 days of growth at 30°C.

### Agrobacterial insertional mutagenesis

*Agrobacterium tumefaciens* bearing a the *T-DNA* plasmid with the *C. neoformans* NAT resistance cassette (McClelland *et al*. 2005) were cultured in 120 ml AMM (0.35% potassium phosphate, 2.6 mM sodium chloride, 2 mM magnesium sulfate, 0.45 mM calcium chloride, 10 μM iron sulfate, 0.05% ammonium sulfate and 0.2% glucose) with 10 μg/ml kanamycin for at least 16 hours at 28°C with shaking. Cells were harvested by centrifugation at 4500xg for 15 min at room temperature and enough bacteria resuspended in 20 ml induction medium (40 mM 2-(N-morpholino)ethanesulfonic acid or MES, pH 5.3, 3% sucrose, 0.5% glycerol) with 10 μg/ml kanamycin and 200 μM acetosyringone to achieve an optical density of 0.15 at 600 nm. This culture was then incubated an additional 6 hours at 28°C and then harvested by centrifugation at 4500xg for 15 min. Finally, the cells were resuspended in 10 ml induction medium and adjusted to an OD_600_ of 1.25 to yield approximately 30 ml.

The *ura5::HAR1* strain was also cultured overnight in YPAD at 30°C with shaking. *C. neoformans* cells were diluted to an OD_600_ of 1.0 and grown for an additional 6 hours at 30°C with shaking. The cells were harvested by centrifugation for 5 min at 2000xg, washed twice with sterile ddH_2_O and resuspended in 10 ml induction media. The volume was adjusted to achieve an OD_600_ of 5.85 with induction medium.

Equal volumes of *A. tumefaciens* and *C. neoformans* were mixed together to yield 500 μl per plate and spread onto an OSMONICS Nylon membrane on induction medium plates with 200 μM acetosyringone and 0.6% agar. Induction plates were incubated upside down (agar down) for 72 hours at room temperature. Membranes were then transferred to YPAD plates containing 75 μg/ml carbenicillin, 100 μg/ml NAT and 200 μM cefotaxime and incubated at 30°C for 48 hours. Once colonies appeared, they were replica plated using sterile velvets onto new plates composed of the same medium and incubated at 30°C for 24 hours. Following this final selection step, the colonies were replica plated onto synthetic complete medium lacking uracil and incubated at 30°C for up to 6 days, checking each day for the appearance of new colonies. The YPAD/NAT/Cefotaxime plates from the previous steps were retained to determine the background level of insertions.

### Determination of insertion sites

Genomic DNA was prepared from the NATR+ and NATR+/Ura+ pools as previously described (Chun and Madhani 2010). 20 μg of genomic DNA was sonicated for 10 min (30 sec on, 1 min rest) at 4°C. DNA was extracted with phenol/chloroform/isoamyl alcohol (25:24:1, Sigma), washed with chloroform and precipitated with 3 volumes ethanol. The extent of the fragmentation was determined by separation on a 0.8% agarose gel.

Linear PCR originating in the *T-DNA*:NAT insertion was performed with Accuprime Taq High Fidelity polymerase (Thermo Fisher) using the JEBPN-Biotin2 primer (Table S6). First strand DNA was then purified over M280 Dynabeads (Thermo Fisher) according to the manufacturer’s instructions. Purified linearized DNA was ligated to the JEBPN-DNA linker (Table S6) using Circligase II (Epicentre) according to the manufacturer’s instructions. Finally, DNA was amplified by nested PCR with JEBPN-SA-II and JEBPN_index-SA-I (Table S6) with various indexes using Accuprime Taq High Fidelity polymerase (Thermo Fisher). Libraries were size selected by non-denaturing PAGE (8% Novex TBE, Thermo Fisher) and extracted from the gel by crushing and elution into 0.3 M sodium acetate overnight at 4°C. The libraries were precipitated in isopropanol and sample quality and quantity was assessed by the Agilent Bioanalyzer High Sensitivity DNA assay and qPCR. Libraries were mixed with a PhiX sample to improve sequence diversity and sequenced on a HiSeq2500 with the JEBPN-SP3 primer (Table S6).

Data were pre-processed for alignment using a custom script as follows: First, reads were filtered for those beginning with at least 6 nt of the *T-DNA* sequence (“TTGTCTAAGCGTCAATTTGTTTACACCACAATATATC”). Second, the adaptor was removed from the 3’ end of the reads (“GTATGCCGTCTTCTGCTTG”). Trimmed reads that were at least 18 nt long were retained for alignment to the genome with bowtie1 (additional parameters: −v2 and −m1) (Langmead *et al*. 2009). Samtools (Li *et al*. 2009) was used to convert, sort and index bam files. Bedgraph files were generated with BEDTools (Quinlan and Hall 2010) and visualized in the Integrative Genomics Viewer (IGV) (Robinson *et al*. 2011). Reads in annotated genes were counted using HTseq-count (Anders *et al*. 2015).

### Targeted mutagenic screening of *PRP43*

A library of *PRP43* alleles was generated by mutagenic PCR (Cadwell and Joyce 2006) of the *PRP43* open reading frame. The mutagenized PRP43 amplicons were combined with the G418 resistance cassette in the pRS316 backbone by in vivo recombination in *S. cerevisiae* (Finnigan and Thorner 2015). Repaired plasmids were retrieved from *S. cerevisiae* by DNA extraction and electroporated in DH10β cells. All *E. coli* transformants were pooled and the plasmid library was prepared using a Qiagen Maxi Prep kit. The mutagenic diversity of the library as assessed by retransformation into *E. coli* followed by miniprep and Sanger sequencing. The plasmid library was then linearized with HindIII (NEB) and introduced into the *ura5::HAR1* screening strain by biolistic transformation.

Strains resistant to both G418 and NAT were then further screened for growth defects at 25°C, 30°C, 34°C and 37°C as well as growth on SC-Ura medium indicative of *HARBINGER* mobilization. The identified alleles were then reconstructed by amplification of the *PRP43* coding sequence from genomic DNA and incorporation into the same plasmid construct in the manner described above. The HindIII linearized plasmid was introduced by biolistic transformation into *C. neoformans* and the allele was confirmed by colony PCR and Sanger sequencing.

### RNA preparation and siRNA Northern blotting

RNA samples were prepared as previously described (Dumesic *et al*. 2013) from log phase YPAD cultures. 30 μg of total RNA were desiccated, re-dissolved in formamide loading dye and separated on a 15% TBE-Urea gel (Novagen) in 1X TBE at 180 V for 80 min. RNA was transferred to Hybond-NX membrane (Amersham) 1X TBE in a Invitrogen Xcell II blot module for 90 min at 20 V. RNA was crosslinked to the membrane in 0.16 M N-(3-Dimethylaminopropyl)-N’-ethylcarbodiimide hydrochloride (Sigma) in 0.13 M 1-methylimidazole (Sigma), pH 8 for 1 hour at 60°C. The membrane was equilibrated in 10 ml Roche Easy Hyb solution at 25°C. Riboprobe against <u>*CNAG_06705*</u> antisense RNA was prepared using the Roche Dig Northern Starter Kit according to the manufacturer’s instructions. 10 μl of the riboprobe was hydrolyzed in 120 μl 0.1 M sodium carbonate + 180 μl 0.1M sodium bicarbonate for 30 min at 60°C then added directly to the Easy Hyb solution. 10 ng/ml of Dig-anti-U6 probe (Table S6) was also added to the hybridization mixture. The probe was hybridized to the membrane overnight at 25°C. The membrane was then washed twice with 6X SSC, 0.1% SDS twice at 37°C for 10 min then once with 2X SSC, 0.1% SDS at 25°C for 10 min. Detection of digoxin on the membrane was performed using the Roche Dig Northern Starter Kit according to the manufacturer’s instructions. Chemiluminescence was detected using an Azure c600.

### RNA-seq and siRNA-seq library preparation

For RNA-seq (all except *rrp6Δ*), 2.5 μg of RNA were treated with DNase as previously described (Zhang *et al*. 2012). 3’ end RNA sequencing libraries were prepared with the Lexogen QuantSeq 3’ mRNA-Seq Library Prep Kit FWD. For RNA from the *rrp6Δ* strain and a corresponding wild type sample, 50 μg of RNA was first selected using the Qiagen Oligotex mRNA mini kit following the manufacturer’s instructions and DNase treated as for QuantSeq samples. Libraries were then prepared using the NEBNext Ultra Directional RNA Library Prep Kit for Illumina following the manufacturer’s instructions. For small RNA-seq, 20 μg of RNA were treated with DNase as previously described (Zhang *et al*. 2012). 3’ end RNA sequencing libraries were prepared with the Lexogen Small RNA-Seq Library Prep Kit.

### Analysis of RNA-seq data

Reads were prepared for alignment by trimming adaptor sequences with Cutadapt (A_10_ for QuantSeq and “TGGAATTCTC” for small RNA-seq) (Martin 2011). Trimmed reads were aligned to the *C. neoformans* genome with either STAR for QuantSeq data (Dobin *et al*. 2013) or Bowtie for small RNA-seq data (Langmead *et al*. 2009). Split reads were ignored when aligning with STAR (--alignIntronMax 1). For small RNA-seq, 2 mismatches were allowed (-v2) and reads aligning to more than one locus were randomly assigned (-M1 --best). Reads aligning to genes and transposable elements were counted using custom scripts using the Python library Pysam and differential expression of mRNAs and siRNAs was determined using DESeq2 (Love *et al*. 2014). The location of transposable elements was determined based on homology with the consensus sequences determined from *C. neoformans var neoformans* (Janbon *et al*. 2010) with custom scripts using BLAST (McGinnis and Madden 2004).

### Mass spectrometry

Tandem immunoprecipitation using C-terminal 2xFLAG and calmodulin binding peptide epitopes followed by mass spectrometry was performed similarly to previously described methods (Dumesic
 *et al*. 2013) from 2 L of *C. neoformans* cultured in YPAD to OD 2.

### Reagent and data availability

Strains are available upon request and are listed in Table S6. Mass spectrometry and processed sequencing data are available as supplemental tables (Tables S1-S5). Raw and processed sequencing data are available from GEO with accession code GSE128009.

## Results

### A colony-level assay to measure the mobilization of a *HARBINGER* DNA transposon

To screen for previously unreported factors involved in silencing transposable elements, we developed a reporter that links uracil prototrophy to mobilization of the *HARBINGER* DNA transposon. *HARBINGER* is suppressed by the RNAi pathway in *C. neoformans* (Wang *et al*. 2010). A copy of *HARBINGER, HAR1* (<u>*CNAG_02711*</u>) was inserted into the second intron of *URA5* (<u>*CNAG_03196*</u>), grossly disrupting the intron and resulting in uracil auxotrophy (Figure 1A). To estimate the rate of *HAR1* transposition, cell populations in which *URA5* is still disrupted by *HARBINGER* were isolated by selection on 5-FOA. The selected cells were then transferred directly to rich media (YPAD) or media lacking uracil, and colony forming units (CFUs) were counted between 2-6 days at 30°C (Figure 1B). Upon deletion of genes encoding a core RNAi factor such as Argonaute (*AGO1*) or the RNA-dependent RNA polymerase (*RDP1*), the number of CFUs on media lacking uracil increased 1,000 to 10,000-fold in comparison to rich media (Figure 1C), consistent with loss of RNAi-based transposition suppression. The *ura5::HAR1* system provides a platform for high-throughput screening for factors involved in *HARBINGER* silencing.

**Figure 1.**
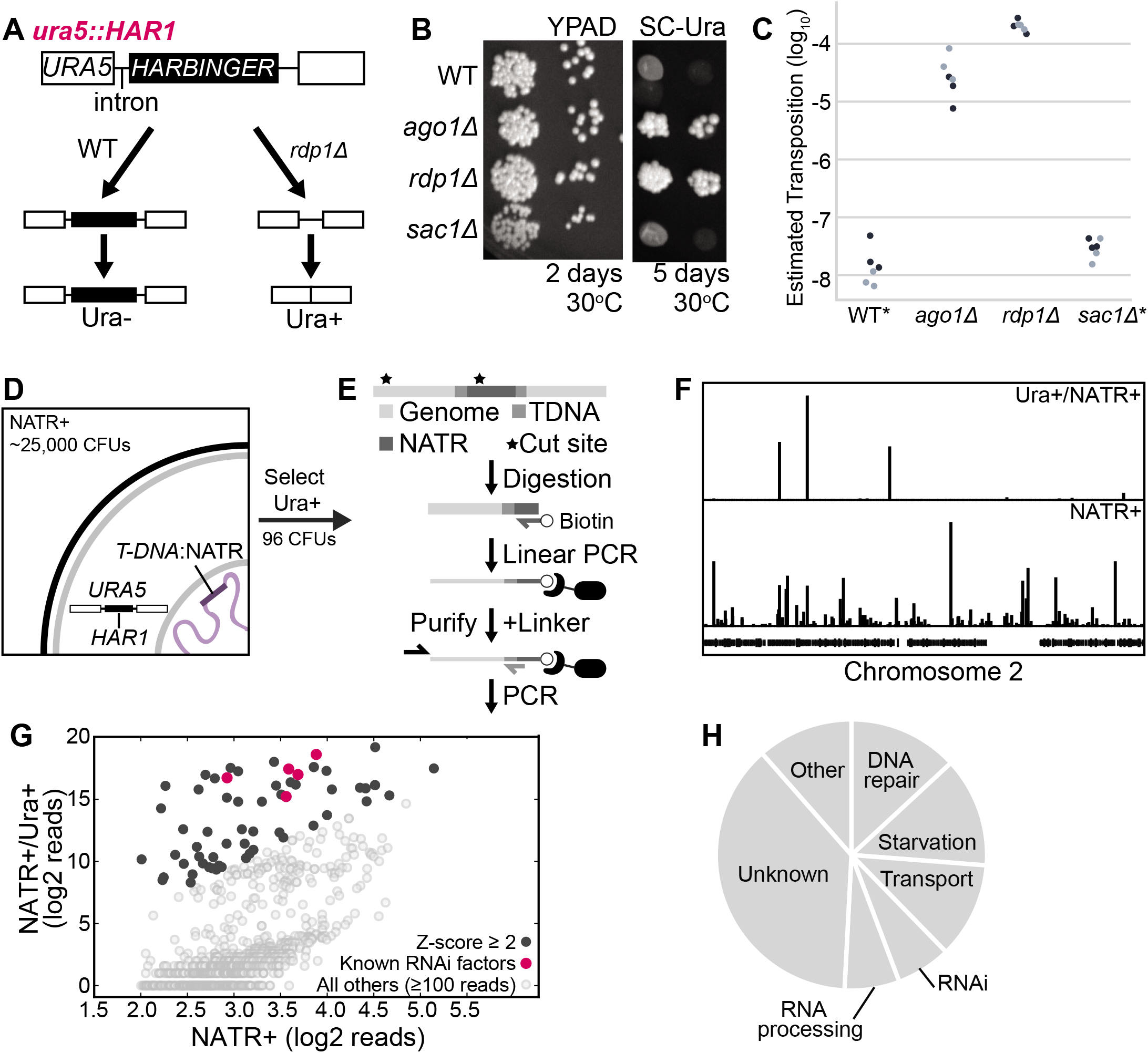
**A**. A copy of the *HARBINGER* transposon (*HAR1*) was inserted into the second intron of *URA5*, resulting in failure to splice and uracil auxotrophy when the RNAi pathway is functional. Upon loss of RNAi (*rdp1Δ*), transposition may occur and some cells are able to synthesize uracil. **B**. Mobilization of *HARBINGER* in the presence of RNAi pathway knockouts. 1/10 dilutions of log phase cultures were spotted on YPAD or SC-Ura and incubated at 30°C for 2 or 5 days. **C**. Quantitation of *HARBINGER* mobilization. CFUs were counted after 2 days (YPAD) or 6 days (SC-Ura) at 30°C. Ura+ CFUs were then normalized to YPAD CFUs. *No colonies in WT and *sac1* genotypes were detected on SC-Ura by the end 6 days, so a maximum estimated transposition rate is indicated. **D**. Schematic of the insertional mutagenesis strategy used for screening. Cells co-cultured with *Agrobacteria* carrying a *T-DNA*:NATR transposable element were selected for resistance to NAT, then for the ability to grow on media lacking uracil. **E**. Sequencing strategy for identifying *T-DNA* insertions. *C. neoformans* genomic DNA was fragmented by sonication and then ssDNA against the insertion site was generated by linear amplification. The biotinylated ssDNA product was purified, a DNA linker of known sequence was added to the 5’ end and the genomic flank was amplified by nested PCR. **F**. Mapping of insertion sites to the *C. neoformans* genome in the NAT resistant uracil prototrophic pool (NATR+/Ura+) versus NAT resistant pool (NATR+). **G**. Quantitative comparison of the number of reads spanning the *T-DNA* boundaries for insertions within annotated genes for NATR+ and NATR+/Ura+ pools. Z-scores were determined from the distribution of the log_2_ ratio of reads from the NATR+/Ura+ pool over reads from the NATR+ pool. **H**. Functional classification of genes with enriched insertions in the *HARBINGER* mobilization screen based on FungiDB and hand-curated annotations.

### Insertional mutagenesis uncovers previously unidentified RNAi factors

We next performed insertional mutagenesis of the *C. neoformans* genome in the *ura5::HAR1* background. A strain of *A. tumefaciens* carrying a nourseothricin (NAT) resistance cassette bounded by *T-DNA* inverted repeat terminal sequences (McClelland *et al*. 2005) was co-cultured with *C. neoformans* resulting in ~25,000 NAT-resistant transformants (Figure 1D). The mutant pool yielded 96 uracil prototroph strains within 2-6 days at 30°C.

T-DNA insertions in all NAT resistant (NATR+) and uracil prototroph (Ura+) strains were identified by performing linear PCR originating in the NATR cassette followed by purification of the biotinylated ssDNA from genomic DNA [Figure 1E, adapted from (Schmidt *et al*. 2007)]. The upstream flanking sequence was then amplified by nested PCR and insertion sites were identified by high throughput sequencing and alignment to the *C. neoformans* genome (Figure 1F). We detected insertions with at least 10 reads in 1804 genes in the NATR+ pool and 752 genes in the Ura+/NATR+ pool. We further refined the list of enriched insertion sites by determining Z-scores from the log_2_ ratio of reads in the Ura+/NATR+ pool to reads in the NATR+ pool (Figure 1G, black, Table S1). The 97 genes with a Z-score of at least 2.0 included four genes encoding known RNAi factors (*AGO1, RDP1, GWO1*, and *QIP1*) (Dumesic *et al*. 2013) and the gene neighboring *RDP1* (Figure 1G, magenta). Interestingly, an insertion in one of the *HARBINGER* loci, *HAR2* (<u>*CNAG_00903*</u>) exhibited two-fold enrichment in the uracil prototroph pool (Table S1), suggesting that decreased expression of *HAR2* might improve the ability of *HAR1* to transpose. The remaining annotated hits primarily occur in factors involved in starvation response, nutrient and small molecule transport, DNA repair and RNA processing (Figure 1H).

To validate the initial hits of the screen, we isolated RNA from 52 available coding sequence knockouts from a *C. neoformans* knockout collection being constructed in our laboratory and detected siRNAs against the endo siRNA-producing locus *<u>CNAG 06705</u>* by small RNA northern analysis (data not shown). Five of the knockouts that exhibited partial or complete loss of siRNAs (Figure 2A) were selected for further study and named *RDE1-5* (Figure 2B; RNAi-DEfective). Knockouts of these five genes also exhibited decreased levels of siRNAs against all three copies of *HARBINGER* (Figure 2C) and dramatically increased transposition of *HAR1* (Figure 2D and Table S2). Finally, RNA-seq analysis of the knockout strains revealed minimal differential transcript expression and no reduction in the expression of RNAi pathway members (Figure 2E and Table S3).

**Figure 2.**
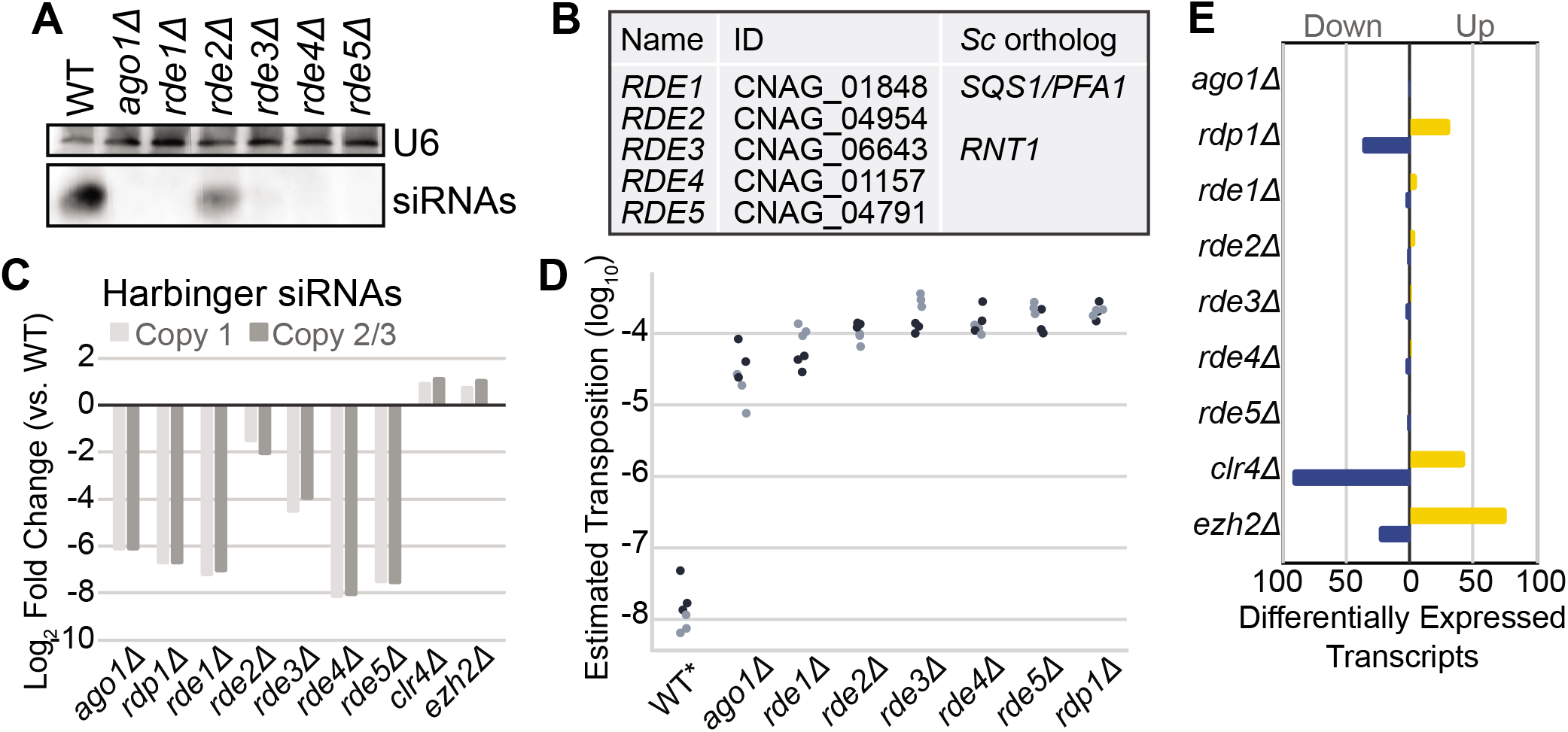
**A**. siRNA Northern analysis of screen hits. Shown are those with a qualitatively apparent loss of siRNAs. Total RNA from each strain was separated by denaturing PAGE, transferred, chemically crosslinked and probed for siRNAs against <u>*CNAG_6705*</u> with a DIG-labeled riboprobe. The loading control, U6 snRNA, was detected with a DIG-labeled DNA oligo against the *C. neoformans* U6 sequence. **B**. Names, gene identifiers and orthologs from *S. cerevisiae* (*Sc*) as determined by PSI blast (Altschul *et al*. 1997). **C**. Quantitation of the fold change in siRNAs against each *HARBINGER* locus (Copy 1: <u>*CNAG_00903*</u>, Copy 2/3: <u>*CNAG_02711*</u> and <u>*CNAG_00549*</u>) as determined by small RNA-seq. Two copies of *HARBINGER* are virtually identical (Copy 2/3) and thus reads map to both with roughly equivalent frequency in our alignment strategy (see methods). **D**. Estimated transposition rate of *HARBINGER* in knockouts of each of the newly discovered factors. Assay conducted as described in Figure 1 legend. **E**. Differential expression of mRNA in *C. neoformans* (QuantSeq). Transcripts were determined to be significantly differentially expressed if they exhibit at least a 2-fold increase (yellow) or decrease (blue) in expression with an adjusted p-value of at most 0.01 (as determined by DESeq2, 2 replicates).

### Mutants of new RNAi pathway members display defects in siRNA accumulation

To determine the extent and specificity of the RNAi defect in the newly discovered RNAi pathway members, we compared the small RNA populations in the knockouts to wild type and knockouts of canonical pathway members by sequencing the total 15-30 nt RNA population in the cell (small RNA-seq). In wild type, we observe that the majority of reads fall within the range of 21-24 nt (Figure 3A-B). Upon loss of a member of the RNAi pathway such as *AGO1*, this peak is no longer discernable (Figure 3A, panel 1, purple) and the proportion of reads between 21-24 nt decreases (Figure 3B). In contrast, when transcriptional silencing of siRNA targets in heterochromatic regions is lost, as in the case of *clr4Δ* (whose gene encodes the sole H3K9 methylase in *C. neoformans*) the 21-24 nt small RNA population increases (Figure 3A, panel 1, yellow). In mutants of each of the factors identified in this screen we observe a different extent of loss of 21-24 nt siRNAs with *rde4Δ* and *rde5Δ* exhibiting the most severe loss and *rde2Δ* exhibiting the least (Figure 3A-B).

**Figure 3.**
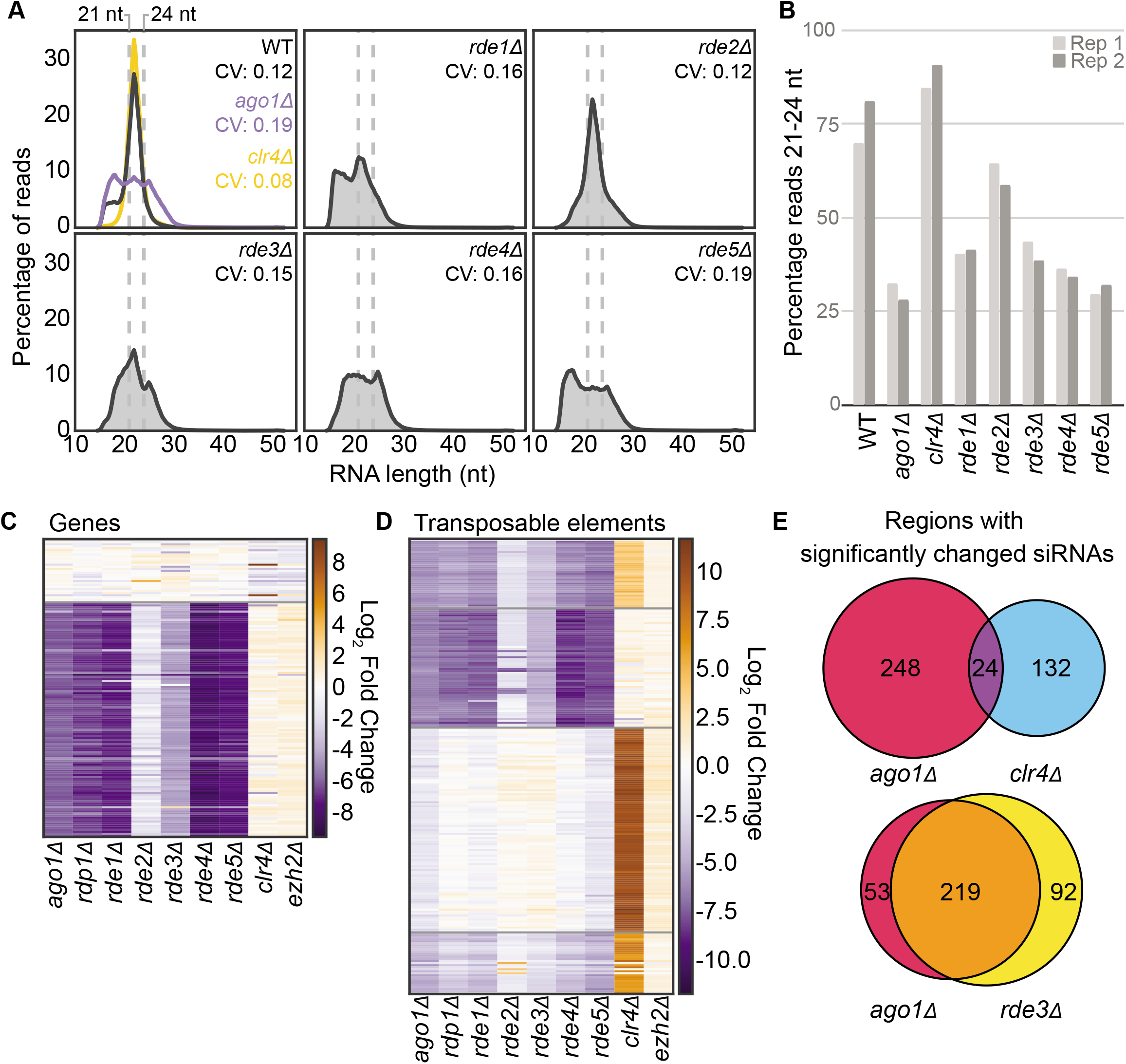
**A**. Distribution of small RNA-seq read sizes and coefficient of variation (CV) in each *RDE* knockout as well as *ago1Δ* and a partial deletion of *clr4Δ* (single replicate shown). **B**. Percentage of small RNA-seq reads between 21-24 nt in length for each knockout compared to wild type. **C**. Fold change of siRNA abundance for annotated genes with at least 100 small RNA-seq reads in both of the wild-type libraries (determined by DESeq2, 2 replicates). Order determined by K-means clustering. **D**. Fold change of siRNA abundance (determined by DESeq2, 2 replicates) against transposable elements and transposable element remnants (see methods for annotation strategy) compared to wild type. **E**. Overlap of transposable elements and genes with significantly changed siRNA populations.

To address whether the observed decrease in siRNAs was specific to certain targets, we also counted small RNA reads antisense to genes (Figure 3C, Table S4) or transposable elements (Figure 3D, Table S4). We observe a similar pattern of siRNA abundance changes in the knockouts of RNAi pathway members and the newly discovered factors (Figure 3C-D), which differs primarily in the magnitude of the small RNA loss. Notably, siRNAs that decrease when RNAi pathway members are lost are largely mutually exclusive with siRNAs that increase when heterochromatic silencing is lost upon deletion of *CLR4* (Figure 3D-E). Additionally, deletion of *EZH2*, the histone 3 lysine 27 methyltransferase component of Polycomb complex (Dumesic *et al*. 2015) has little effect on the small RNA population (Figure 3C-D). Taken together, these results indicate that the factors identified in our screen are required for the biosynthesis of endo-siRNAs in *C. neoformans*, while heterochromatin formation antagonizes siRNA production, presumably by limiting expression of the transcriptions that serve as templates for siRNA production.

### Nucleolar protein homologs required for endo-siRNA production

To expand our understanding of the function and localization of the newly discovered RNAi factors, we performed tandem affinity purification followed by label-free mass spectrometry analysis on each Rde factor using a C-terminal epitope tag [Calmodulin binding protein (CBP), 2xFLAG]. Rde1, which contains a G-patch domain thought to interact with DEXD-box helicases (Figure 4A), primarily associates with ribosomal proteins, factors involved in translation and nucleolar proteins (Figure 4B, Table S5). Rde1 associates with the essential DEXD-Box helicase Prp43, which is both a ribosome biogenesis and pre-mRNA splicing factor. The *S. cerevisiae* ortholog of Rde1, Sqs1/Pfa1, also associates with Prp43 and is implicated in ribosome biogenesis (Lebaron *et al*. 2009; Pandit *et al*. 2009; Pertschy *et al*. 2009). Notably, Rde1 also associates with Ago1 and Rde2 as well as Rnh1, an RNase H that cleaves RNA-DNA duplexes. Rde2 associates with Rde1 as well as Rnh1 and a few other overlapping proteins; however, Prp43 and Ago1 were not detected in the Rde2-CBP-2XFLAG purification sample (Figure 4C).

**Figure 4.**
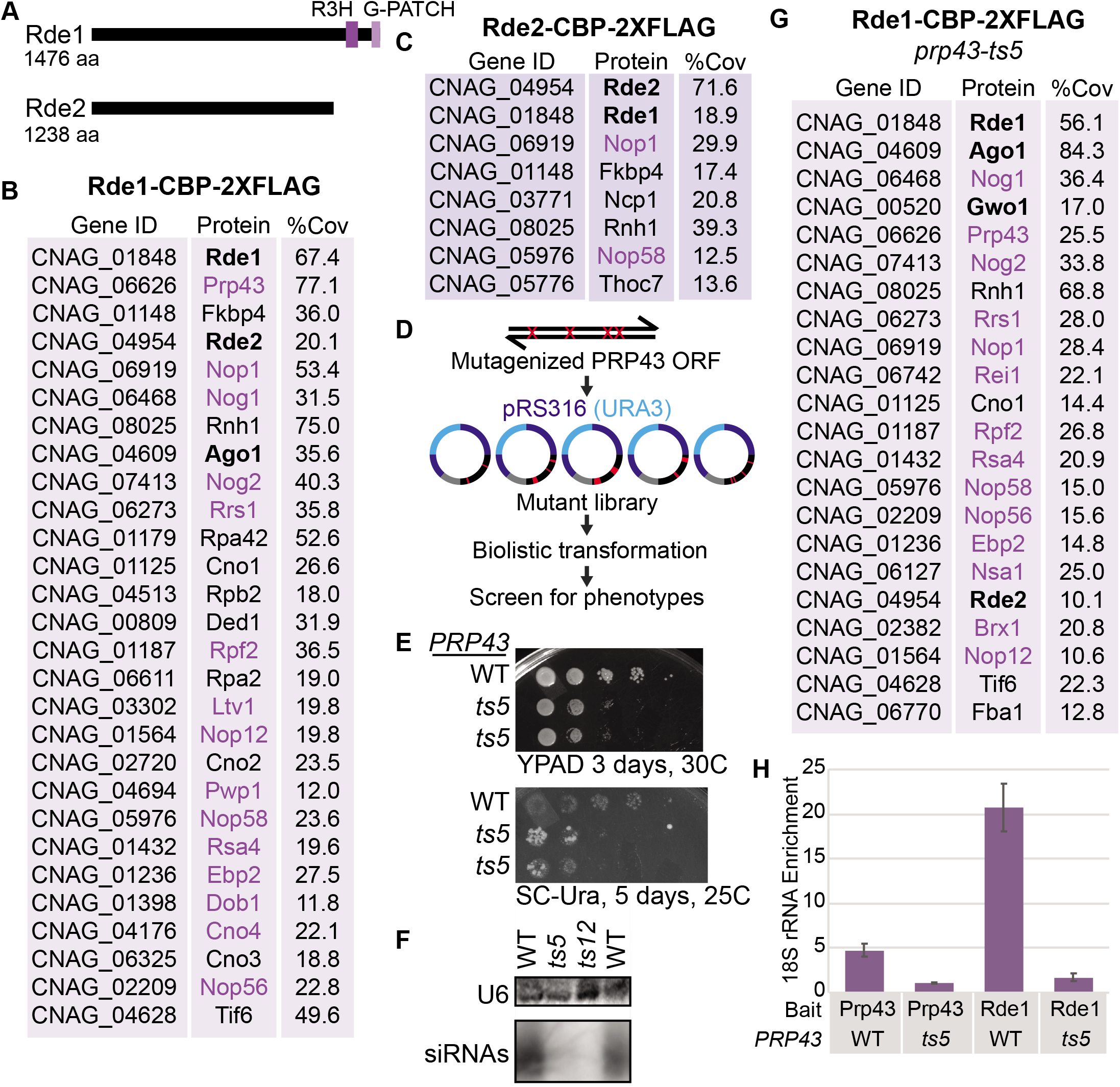
**A**. Predicted domain structures of Rde1 and Rde2 determined by PSI-BLAST with an e-value of at most 1X10^−5^. **B**. Proteins associated with Rde1 as determined by tandem IP-MS, named based on current *C. neoformans* annotation (FungiDB) or *S. cerevisiae* homolog. Three factors had no clear homolog, so we refer to them as putative <u>*C*</u>. *neoformans* <u>N</u>ucle<u>O</u>lar protein 1-3 (Cno1, Cno2 and Cno3). Proteins in bold were identified in the screen. Proteins in purple are predicted nucleolar proteins, typically involved in rRNA processing and ribosome biogenesis. Percent coverage is the average between two replicates. Common contaminants and proteins with less than 10% coverage are excluded and proteins are in order of descending total spectral counts. Only proteins with at least 30 spectral counts are shown, but the remainder can be found in Table S5. **C**. Proteins associated with Rde2 (see B), data from only one replicate. **D**. Mutagenic library strategy for *PRP43*. **E**. Growth phenotype of wild type and *prp43-ts5 C. neoformans* strains bearing the *ura5::HAR1* insertion on rich media and media lacking uracil. **F**. Loss of siRNAs against <u>CNAG 06705</u> as determined by Northern analysis (see Fig. 2A). **G**. RT-qPCR of 18S rRNA associated with Rde1 (native FLAG affinity purification). Data are from three technical replicates and two biological replicates.

To determine whether Prp43 function is important for endo-siRNA biogenesis, we generated a randomly mutagenized library of *PRP43* alleles and screened for mutants that increased transposition of *HAR1* using the *ura5::HAR1* system (Figure 4D). We identified two alleles of *PRP43*, termed *prp43-ts5* (N207H, Y214C, K277R, T290A, H515R, R620Q, K778R) and *prp43-ts12* (F708S), that result in increased transposition of *HARBINGER* as well as a severe growth defect at 37°C (data not shown). Reconstructions of these alleles also exhibited increased transposition of *HAR1* (Figure 4E, *prp43-ts12* not shown) and loss of siRNA production (Figure 4F). In the presence of *prp43-ts5*, Rde1 appears to more strongly associate with Ago1 and also associates with the P-body localized RNAi factor, Gwo1. Additionally, RT-qPCR reveals that enrichment for the 18S ribosomal RNA by both Rde1 and Prp43 is reduced in the *prp43-ts5* background (Figure 4H).

### RNA processing factors link RNAi with RNA surveillance

The remaining RNAi factors contain domains that suggest they are involved in mRNA processing and RNA surveillance (Figure 5A). Rde3 contains an RNase III domain, but no PAZ domain, suggesting that it can cleave double stranded RNA (dsRNA) but is not a canonical Dicer enzyme. Rde4 contains a terminal-nucleotidyl transferase domain commonly found in terminal-uridylyl transferases and polyA polymerases, while Rde5 has no strong domain predictions.

**Figure 5.**
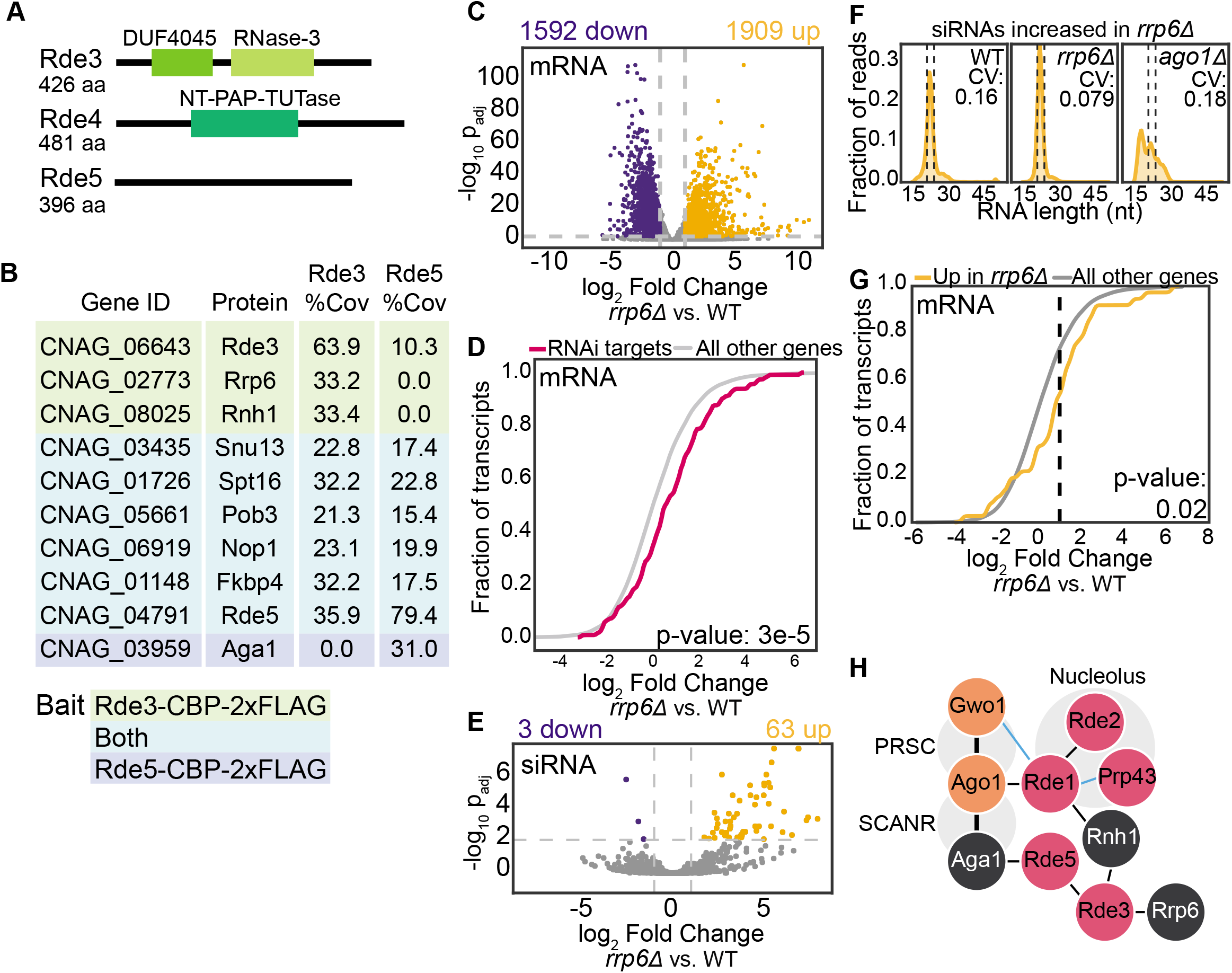
**A**. Predicted domain structures of Rde3, Rde4 and Rde5 determined by PSI-BLAST (Altschul *et al*. 1997) with an e-value of at most 1X10^−5^. **B**. Proteins associated with Rde3 and Rde5 as determined by tandem affinity purification and mass spectrometry. Proteins highlighted in green were found associated only with Rde3; those highlighted in teal were associated with both Rde3 and Rde5 and Aga1 (in blue) was associated only with Rde5. Percent coverage is the average between two replicates. Common contaminants and proteins with less than 10% coverage are excluded and proteins are in order of ascending total spectral counts. **C**. Differential expression of polyA mRNA in the *rrp6Δ* strain. Colored points indicate transcripts with at least a 2-fold increase (yellow) or decrease (blue) in expression with an adjusted p-value of at most 0.01 (DE-Seq2). **D**. Fold change of transcripts that template endo-siRNA production (Dumesic *et al*. 2013) compared with the rest of the transcriptome. P-value determined by Mann-Whitney U test. **E**. Differential abundance of small RNAs in the *rrp6Δ* strain. See panel **C** for color-code. **F**. Small RNA size distribution and CV of the population significantly increased in the *rrp6Δ* strain. Dashed lines indicate 21-24 nt region. **G**. Fold change of transcripts targeted by siRNAs that are differentially increased in *rrp6Δ*. P-value determined by Mann-Whitney U test. **H**. Physical associations of new RNAi factors described in this study (pink) with known RNAi factors (orange) and other RNA processing factors (white). Thin lines are interactions from IP-MS data only. Thick lines indicate interactions confirmed by co-IP and/or yeast 2-hybrid (Dumesic *et al*. 2013). Blue lines indicate interactions that are affected by the *prp43-ts5* allele.

In contrast with Rde1-2, Rde3-5 largely do not associate with rRNA processing and nucleolar proteins. Rde3, the putative RNase III, and Rde5 co-associate with each other and a variety of other nuclear proteins (Figure 5B). Rde3 associates with RNA quality control enzymes, such as Rnh1 and Rrp6, a component of the nuclear exosome. Both Rde3 and Rde5 associate with homologs of Spt16 and Pob3, members of the FACT complex involved in chromatin remodeling (Lejeune *et al*. 2007), as well as a homolog of Nop1, a nucleolar protein, and Fkbp4. Interestingly, Rde5 associates with Aga1, which is a known binding partner of Ago1 (Dumesic *et al*. 2013). Rde4, the putative terminal nucleotidyltransferase, did not co-purify with any proteins aside from common contaminants (Table S5).

### Impact on RNAi target transcript levels and siRNA abundance by the nuclear exosome

Finally, to investigate the connection between RNAi and nuclear RNA surveillance and quality control, we performed RNA-seq and small RNA-seq in a strain lacking the nuclear exosome component, Rrp6. Consistent with the known role of Rrp6 in other systems, deletion of *RRP6* results in substantial changes in the mature RNA population (Figure 5C). While many transcripts are significantly more abundant in the *rrp6Δ* strain (Figure 5C), targets of the RNAi pathway exhibit a greater increase in abundance (Figure 5D), suggesting they are turned over at a somewhat higher rate by the nuclear exosome than transcripts on average.

Additionally, a subset of small RNAs increase in abundance in *rrp6Δ* compared with wild type (Figure 5E). However, these small RNAs could just be degradation products of suboptimal transcripts that are normally degraded by the nuclear exosome. The size distribution of the small RNAs increased in *rrp6Δ* indicates that these are canonical 21-24 nt siRNAs (Figure 5F). The small RNAs in these regions are also dependent on *AGO1*, indicating that they are normally produced by the canonical RNAi pathway (Figure 5F). Finally, the mRNA targets of these small RNAs exhibit a modestly significant differential increase in expression in *rrp6Δ* compared with the general mRNA population (Figure 5G). Together with the result that the RNase III Rde3 associates with Rrp6 and other RNA surveillance factors, our findings are consistent with a competition relationship between nuclear RNA surveillance and the production of endo-siRNAs in *C. neoformans*.

## Discussion

In this study, we describe a genetic screen aimed at identifying novel factors involved in transposon mobilization in *C. neoformans*. We report the impact of deletion alleles of five genes identified in the screen, *RDE1-5*, on global endo-siRNA levels, global RNA levels, and transposon mobilization. We tagged each gene and performed tandem affinity purification and mass spectrometry experiments to investigate protein-protein interactions. We also describe two conditional alleles of *C. neoformans PRP43* that impact endo-siRNA levels and transposon mobilization.

Two of these factors, Rde1 (a homolog of the *S. cerevisiae* nucleolar protein Sqs1/Pfa1) and Rde2, associate with one another. Rde1 associates with factors homologous to *S. cerevisiae* proteins that localize to the nucleolus. Additionally, we find that two different mutants of the nuclear/nucleolar helicase Prp43 display reduced siRNA production, further implicating this rRNA processing machinery in siRNA biogenesis. Given that Prp43 also disassembles stalled and post-catalytic spliceosomes and that stalled spliceosomes can serve to trigger siRNA production in *C. neoformans* (Dumesic et al., 2013), it is possible that Prp43 is released from the nucleolus in these mutants enabling it to disassemble otherwise stalled spliceosomes in the nucleoplasm, thereby inhibiting RNAi. Consistent with this view, we find that the mutant Prp43 alleles display decreased association with 18S RNA and altered protein interactions. Alternatively, the apparently increased association of Rde1 with Ago1 in the *prp43-ts5* strain may point to sequestration of Ago1, perhaps via relocalization in the nucleolus, as an alternative possible mechanism of mutant action. While further work will be required to understand the underlying mechanisms, our results point to a connection between RNAi and the nucleolus.

Rde4, a predicted terminal nucleotidyl transferase that resembles polyA polymerases and terminal-uridylyl transferases (TUTases), is also required for siRNA biogenesis. In some other systems, TUTases have been reported to dampen the effectiveness of the RNAi pathway by inactivating small RNAs and miRNAs (Pisacane and Halic 2017). However, we do not observe any evidence for this in the *C. neoformans* system. We are unable to detect oligoA/U tails on small RNAs in our data and Rde4 promotes rather than inhibits siRNA biogenesis. Rde4 may be responsible for marking for RNAi suboptimal transcripts detected by RNA surveillance that would otherwise target them for degradation by the nuclear exosome (Lim *et al*. 2014). Additionally, the recent finding that LINE-1 elements are modified by TUT4/TUT7 uridylyltransferases to impede mobilization (Warkocki *et al*. 2018) suggests that these modifications may also be protective against proliferation of transposable elements.

Finally, we observe that Rde3, an RNase III related to *S. cerevisiae* Rnt1, and Rde5, a protein of unknown function, associate with one another and that Rde3 associates with homologs of the RNA surveillance factors Rrp6 and Rnh1. In *S. pombe*, a RNAi system mediates formation of heterochromatin in pericentromeric regions presumably via production of dsRNA against non-coding RNA transcripts by an RNA-dependent RNA polymerase (Volpe *et al*. 2002) and subsequent production of siRNAs that direct Ago1. Formation of heterochromatin in *S. pombe* is dependent on the RNAi pathway as well as other factors including Rrp6 and the RNA polymerase II pausing and termination factor Seb1 (Reyes-Turcu *et al*. 2011; Parsa *et al*. 2018). However, disruption of the these factors does not affect siRNA abundance (Bühler *et al*. 2007; Marina *et al*. 2013) indicating that they function in parallel with the RNAi system to maintain heterochromatin.

Our findings indicate that the *C. neoformans* heterochromatin pathway is not required for RNAi as it is in *S. pombe*. In *C. neoformans*, heterochromatin likely silences transposable element transcription at centromeres which in this species limits the production of transcripts that template endo-siRNA production. This model would explain our finding that global endo-siRNAs increase rather than decrease in abundance in cells lacking H3K9me. As RNAi does not generally impact transcript levels in *C. neoformans*, it likely acts at another level such as nuclear export or translation. This two-level mechanism may enable more stringent transposon silencing. Deletion of *RRP6* does not affect the abundance of siRNAs, suggesting that RNAi and exosome-mediated surveillance act in parallel to inactivate target transcripts in *C. neoformans*, providing a third potential layer of transposon suppression.

Based on our analysis of transposition phenotypes of the five genes described here, our screen may have been biased towards identifying strong effects on DNA transposon mobilization. Moreover, as *HAR1* is present in a non-heterochromatic region, factors selectively involved in silencing of retrotransposons, all of which lie in heterochromatic centromeric regions in *C. neoformans*, would have been missed. Screens with a broader dynamic range and those aimed at centromeric elements are thus likely to reveal additional factors that limit transposon mobilization.

## Acknowledgements

We would like to thank all members of the Madhani lab for helpful discussions. We thank Dr. Sandra Catania for construction of the CM1926 strain and Nguyen Nguyen for technical support. We also thank Eric Chow and the UCSF Center for Advanced Technology for assistance with sequencing and sample analysis. We thank anonymous reviewers for critical comments on the manuscript. Supported by NIH grants R01 GM71801 to H.D.M., P41 GM103533 to J.R., and R01 GM120507 to J.J.L. J.E.B was supported by postdoctoral fellowship 127531-PF-15-050-01-R from the American Cancer Society. H.D.M. is a Chan-Zuckerberg Biohub Investigator.

## Tables

**Table S1**: Counts of reads spanning T-DNA insertion sites for all insertions within genes with Z-score analysis. Related to Figure 1.

**Table S2**: Transposition assay CFU counts and estimated transposition rates for wild type and mutant *C. neoformans ura5::HAR1* strains. Related to Figure 2.

**Table S3**: Read counts from QuantSeq, expression changes and significance as determined by DESeq2. Related to Figure 2.

**Table S4**: Read counts from siRNA sequencing, expression changes and significance as determined by DESeq2 and read size distributions. Related to Figure 3.

**Table S5**: Mass spectrometry results from Rde1-5 CBP-2xFLAG affinity purifications. Related to Figure 4–5.

**Table S6**: DNA oligomers and *C. neoformans* strains used in this study.

